# Spatial Release from Informational Masking: Evidence from Functional Near Infrared Spectroscopy

**DOI:** 10.1101/357525

**Authors:** Min Zhang, Antje Ihlefeld

## Abstract

Informational masking (IM) can greatly reduce speech intelligibility, but the neural mechanisms underlying IM are not understood. Binaural differences between target and masker can improve speech perception. In general, improvement in masked speech intelligibility due to provision of spatial cues is called spatial release from masking. Here, we focused on an aspect of spatial release from masking, specifically, the role of spatial attention. We hypothesized that in a situation with IM background sound 1) attention to speech recruits lateral frontal cortex (LFCx), and 2) LFCx activity varies with direction of spatial attention. Using functional near infrared spectroscopy (fNIRS), we assessed LFCx activity bilaterally in normal-hearing listeners. In experiment 1, two talkers were simultaneously presented. Listeners either attended to the target talker (speech task) or they listened passively to an unintelligible, scrambled version of the acoustic mixture (control task). Target and masker differed in pitch and interaural time difference (ITD). Relative to the passive control, LFCx activity increased during attentive listening. Experiment 2 measured how LFCx activity varied with ITD, by testing listeners on the speech task in experiment 1, except that talkers either were spatially separated by ITD or co-located. Results show that directing of auditory attention activates LFCx bilaterally. Moreover, right LFCx is recruited more strongly in the spatially separated as compared with co-located configurations. Findings hint that LFCx function contributes to spatial release from masking in situations with IM.

## INTRODUCTION

In everyday life, background speech often interferes with recognition of target speech. At least two forms of masking contribute to this reduced intelligibility, referred to as *energetic* and *informational* masking (EM and IM, Brungart, 2001; Freyman et al. 2001; Mattys et al. 2009; Jones and Litovsky, 2011). EM occurs when sound sources have energy at the same time and frequency (e.g., Brungart et al. 2006). IM broadly characterizes situations when target and background sources are perceptually similar to each other or when the listener is uncertain about what target features to listen for in an acoustic mixture (for a recent review, see Kidd and Colburn, 2017). IM is thought to be a major factor limiting performance of hearing aid and cochlear implant devices (Shinn-Cunningham and Best, 2008; Marrone et al., 2008; Xia et al., 2017). However, the neural mechanisms underlying IM are not understood. The current study explores cortical processing of speech detection and identification in IM.

In EM-dominated tasks, computational models based on the output of the auditory nerve can closely capture speech identification performance (review: Goldsworthy and Greenberg, 2004). Consistent with this interpretation, subcortical responses encode the fidelity by which a listener processes speech in EM noise (Anderson and Kraus, 2010). However, peripheral models fail to account for speech intelligibility in IM-dominated tasks (e.g., Cooke et al., 2008), suggesting that performance in IM is limited at least partially by mechanisms of the central nervous system.

In IM-dominated tasks, previous behavioral studies are consistent with the idea that in order to understand a masked target voice, listeners need to segregate short-term speech segments from the acoustic mixture, stream these brief segments across time to form a perceptual object and selectively attend to those perceptual features of the target object that distinguish the target talker from competing sound (Jones et al., 1999; Cusack et al., 2004; Ihlefeld and Shinn-Cunningham, 2008a). Previous work suggests that common onsets and harmonicity determine how short-term segments form (Darwin and Hukin, 1998; Micheyl et al., 2010). Differences in higher order perceptual features, including spatial direction and pitch, then allow listeners to link these short-term segments across time to form auditory objects (Darwin and Hukin, 2000; Brungart and Simpson, 2002; Darwin et al., 2003), enabling the listener to selectively attend to a target speaker and ignore the masker (Carlyon 2004; Shinn-Cunningham, 2008; Ihlefeld and Shinn-Cunningham, 2008b).

Rejection of competing auditory streams correlates with behavioral measures of short-term working memory (Conway et al., 2001). This raises the possibility that central regions linked to auditory short-term memory tasks are recruited in situations with IM. To test this prediction, here, we conducted two experiments to characterize blood oxygenation level dependent (BOLD) correlates of cortical responses while normal hearing (NH) subjects listened, either actively or passively, to speech in IM background sound. Recent work in NH listeners demonstrates that auditory short-term memory tasks can alter BOLD signals bilaterally in two areas of lateral frontal cortex (LFCx): 1) the transverse gyrus intersecting precentral sulcus (tgPCS) and 2) the caudal inferior frontal sulcus (cIFS; Michalka et al., 2015; Noyce et al., 2017). Here, we extend this work using functional near infrared spectroscopy (fNIRS) to record BOLD signals at these four regions of interest (ROIs).

In two experiments, we tested rapid-serial auditory presentation stimuli adapted from previous work by Michalka and collagues (2015). Our goal was to examine how direction of auditory attention alters the BOLD responses in LFCx in a situation with IM, as assessed with fNIRS. In experiment 1, NH listeners were asked to detect keywords in a target message on the left side while a background talker producing IM was simultaneously presented on the right. In a control condition, participants listened passively to an unintelligible, acoustically scrambled version of the same stimuli. We hypothesized that unlike in passive listening, when listeners actively tried to hear out speech in IM background sound this would recruit LFCx.

We further hypothesized that interactions between spatially directed auditory attention and LFCx activity would arise. An extensive literature documents that speech intelligibility improves and IM is released, when competing talkers are spatially separated as opposed to being co-located, a phenomenon referred to as spatial release from masking (e.g., Carhart et al, 1967; Darwin and Hukin, 1997; Kidd et al., 2010; Glyde et al., 2013). Using similar speech stimuli as in experiment 1, we looked whether the mechanisms underlying spatial release from IM recruit LFCx, by comparing LFCx BOLD responses in the spatially separated configuration from experiment 1 versus a co-located configuration of the same stimuli. We reasoned that a stronger BOLD response in the spatially separated versus co-located configurations would support the view that spatial attention under IM activates LFCx. In contrast, a stronger LFCx response in the co-located configuration would suggest that LFCx does not encode the direction of spatial auditory attention.

## PARTICIPANTS

A total of 29 listeners (ages 19 to 25, 9 females) participated in the study and were paid for their time, with 14 participants in experiment 1 and 15 participants in experiment 2. All listeners were native speakers of English, righthanded, and had normal audiometric pure-tone detection thresholds as assessed through standard audiometric testing at all octave frequencies from 250 Hz to 8 kHz. At each tested frequency, tone detection thresholds did not differ by more than 10 dB across ears, and all thresholds were 20 dB HL or better. All listeners gave written informed consent to participate in the study. All testing was administered according to the guidelines of the Institutional Review Board of the New Jersey Institute of Technology.

## METHODS

### Recording Setup

Each listener completed one session of behavioral testing while we simultaneously recorded bilateral hemodynamic responses over the listener’s left and right dorsal and ventral LFCx. The listener was seated approximately 0.8 m away from a computer screen with test instructions (Lenovo ThinkPad T440P), inside a testing suite with a moderately quiet background sound level of less than 44 dBA. The listener held a wireless response interface in the lap (Microsoft Xbox 360 Wireless Controller) and wore insert earphones (Etymotic Research ER-2) for delivery of sound stimuli. The setup is shown in Figure 1A.

**Figure 1.**
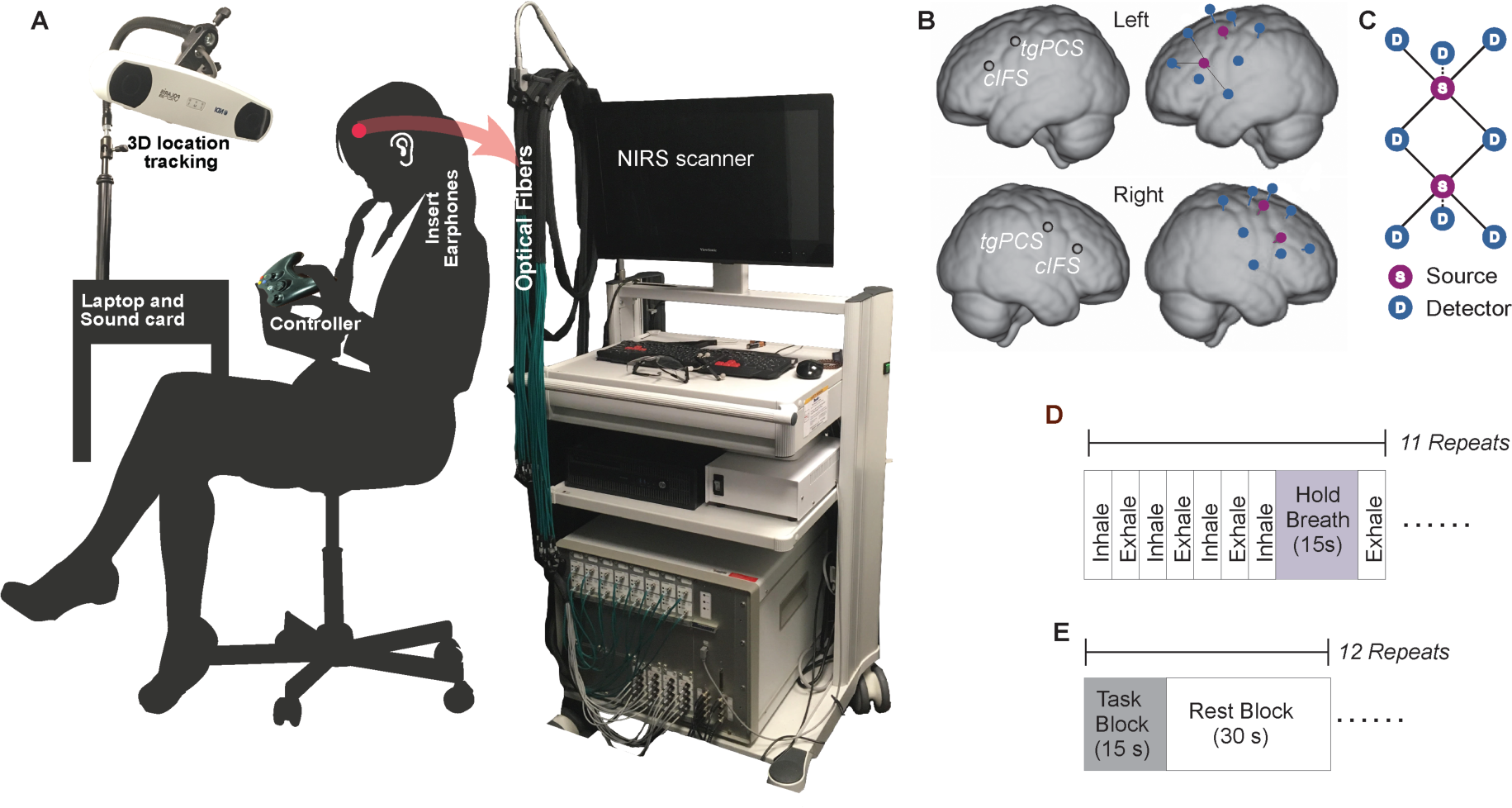
A) Experimental apparatus and setup. B) ROIs and optode placement for a representative listener. Blue circles show placements of detector optodes, red circles of source optodes. C) fNIRS optical probes design with deep neurovascular (solid line) and shallow nuisance (dotted line) channels. S: source. D: detector. D) Block design, Controlled breathing task E) Block design, Auditory task.

A camera-based 3D-location tracking and pointer tool system (Brainsight 2.0 software and hardware by Rogue Research Inc., Canada) allowed the experimenter to record four coordinates on the listener’s head: nasion, inion, and bilateral preauricular points. Following the standard Montreal Neurological Institute (MNI) ICBM-152 brain atlas (Talairach and Tournoux, 1988), these four landmark coordinates were then used as reference for locating the four regions of interest (ROIs, locations illustrated in Fig. 1B). Infrared optodes were placed on the listener’s head directly above the four ROIs, specifically, the left tgPCS, left cIFS, right tgPCS, and right cIFS. A custom-built head cap, fitted to the listener’s head via adjustable straps, embedded the optodes and held them in place.

Acoustic stimuli were generated in Matlab (Release R2016a, The Mathworks, Inc., Natick, MA, USA), D/A converted with a sound card (Emotiva Stealth DC-1; 16 bit resolution, 44.1 kHz sampling frequency) and presented over the insert earphones. This acoustic setup was calibrated with a 2-cc coupler, 1/2” pressure-field microphone and a sound level meter (Bruel&Kjaer 2250-G4).

Using a total of 4 source optodes and 16 detector optodes, a continuous-wave diffuse-optical NIRS system (CW6; TechEn Inc., Milford, MA) simultaneously recorded light absorption at two different wavelengths, 690 and 830 nm, with a sampling frequency of 50 Hz. Sound delivery and optical recordings were synchronized via trigger pulse with a precision of 20 ms. Using a time-multiplexing algorithm developed by Huppert and colleagues (2009), multiple source optodes were paired with multiple detector optodes. A subset of all potential combinations of optode-detector pairs was interpreted as response channels and further analyzed. Specifically, on both sides of the head, we combined one optical source and four detectors into one probe set according to the channel geometry shown in Figure 1C. On each side of the head, we had 2 probe sets placed directly above cIFS and tgPCS on the scalp. Within each source-detector channel, the distance between source and detector determined the depth of the light path relative to the surface of the skull (review: Ferrari and Quaresima, 2012). To enable us to partial out the combined effects of nuisance signals such as cardiac rhythm, respiratory induced change, and blood pressure variations from the desired hemodynamic response driven neural events in cortex, we used two recording depths. Deep channels, used to estimate the neurovascular response of cortical tissue between 0.5 to 1 cm below the surface of the skull, had a 3 cm source-detector distance (solid lines in Fig. 1C), whereas shallow channels, used to estimate physiological noise, had a source-detector distance of 1.5 cm (dotted line in Fig. 1C). At each of the four ROIs, we recorded with four concentrically arranged deep channels and one shallow channel and averaged the traces of the four deep channels, to improve the noise floor. As a result, for each ROI, we obtained one deep trace, which we interpreted as neurovascular activity, and one shallow trace, which we interpreted as nuisance activity.

### Controlled Breathing Task

Variability in skull thickness, skin pigmentation and other idiosyncratic factors can adversely affect recording quality with fNIRS (Yoshitani et al., 2007; Bickler et al., 2013). As a control for reducing group variance and to monitor recording quality, listeners initially performed a non-auditory task, illustrated in Figure 1D. This non-auditory task consisted of 11 blocks of controlled breathing (Thomason et al., 2007). During each of these blocks, visuals on the screen instructed listeners to 1) “Inhale” via a gradually expanding green circle, or 2) “Exhale” via a shrinking green circle, or 3) “Hold breath” via a countdown on the screen. Using this controlled breathing method, listeners were instructed to follow a sequence of inhaling for 5 s, followed by exhaling for 5 s, for a total of 30 s. At the end of this sequence, listeners were instructed to inhale for 5 s and then hold their breath for 15 s. Our criterion for robust recording quality was that for each listener, breath holding needed to induce a significant change in the hemodynamic response at all ROIs (analysis technique and statistical tests described below), otherwise that listener’s data would have been excluded from further analysis. Moreover, we used the overall activation strength of the hemodynamic response during breath holding for normalizing the performance in the auditory tasks (details described below).

### Auditory Tasks

Following the controlled breathing task, listeners performed experiment 1, consisting of 24 blocks of behavioral testing with their eyes closed. Each listener completed 12 consecutive blocks of an active and 12 consecutive blocks of a passive listening task, with task order (active versus passive) counter-balanced across listeners. In each block, two competing auditory streams of 15 s duration each were presented simultaneously. In the active listening task, we presented intelligible speech utterances, whereas in the passive listening task, we presented unintelligible scrambled speech. Figure 2 shows a schematic of the paradigm (A) and spectrograms for two representative stimuli (B).

**Figure 2.**
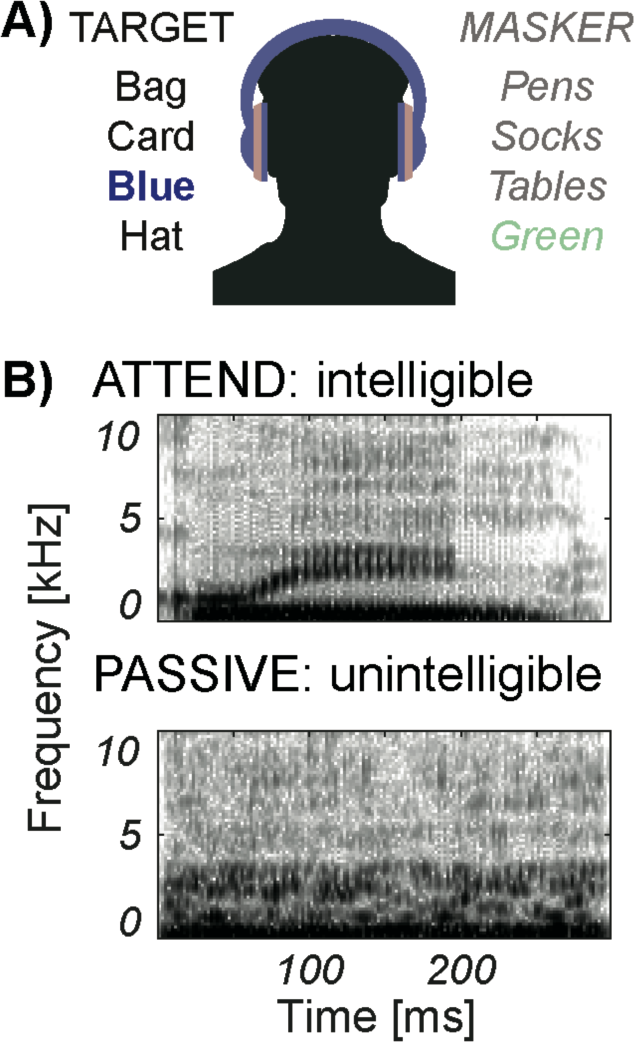
A) Speech paradigm. B) Spectrograms of the word “green.” Unprocessed speech in the ATTEND condition (top) and scrambled speech in the PASSIVE condition (bottom).

In experiment 1, the target stream was always presented with a left-leading interaural time difference (ITD) of 500 μs, while the concurrent masker stream was presented with a right-leading ITD of 500 μs (spatially separated configuration). In experiment 2, we also tested a spatially co-located configuration, where both the target and the masker had 0 μs ITD. In experiment 1, the broadband root means square values of the stimuli were equated at 59 dBA, then randomly roved from 53 to 65 dBA, resulting in broadband signal-to-noise ratios from −6 to 6 dB, so that listeners could not rely on level cues to detect the target. In order to remove level cues entirely, giving spatial cues even more potential strength for helping the listener attend to the target, in experiment 2, we made the target and masker equally loud. In experiment 2, both target and masker were presented at 59 dBA. Unfortunately, due to a programming error, listeners’ responses were inaccurately recorded during the auditory tasks of experiments 1 and 2 and are thus not reported here. During pilot testing with the tested stimulus parameters (not shown here), speech detection performance was 90% correct or better across all conditions.

In the active task, stimuli consisted of two concurrent rapid serial streams of spoken words. Speech utterances were chosen from a closed-set corpus (Kidd et al. 2008). There were sixteen possible words, consisting of the colors <red, white, blue, and green> and the objects <hats, bags, card, chairs, desks, gloves, pens, shoes, socks, spoons, tables, and toys>. Those words were recorded from two male talkers, spoken in isolation. The target talker had an average pitch of 115 Hz versus 144 Hz for the masker talker. Using synchronized overlap-add with fixed-synthesis (Hejna and Musicus, 1991), all original utterances were time-scaled to make each word last 300 ms. Words from both the target and masker talkers were simultaneously presented, in random order with replacement. Specifically, target and masker streams each consisted of 25 words with 300 ms of silence between consecutive words (total duration 15 s).

To familiarize the listener with the target voice, at the beginning of each active block, we presented the target voice speaking the sentence “Bob found five small cards” at 59 dBA and instructed the listeners to remember this voice. Listeners were further instructed to press the right trigger button on the handheld response interface each time the target talker to their left side uttered any of the four *color* words, while ignoring all other words from both the target and the masker. A random number (between three and five) of color words in the target voice would appear during each block. No response feedback was provided to the listener.

In the passive task, we simultaneously presented two streams of concatenated scrambled speech tokens that were processed to be unintelligible. Stimuli in the passive task were derived from the stimuli in the active task. Specifically, using an algorithm by Ellis (2010) unprocessed speech tokens were time-windowed into snippets of 25 ms duration, with 50 *%* temporal overlap between consecutive time-steps. Using a bank of 64 GammaTone filters with center frequencies that were spaced linearly along the human Equivalent Rectangular Bandwidth scale (ERB, Patterson and Holdsworth, 1996) and that had bandwidths of 1.5 ERB, the time-windowed snippets were bandpass filtered. Within each of the 64 frequency bands, the bandpass-filtered time-windowed snippets were permutated with a Gaussian probability distribution over a radius of 250 ms, and added back together, constructing scrambled tokens of speech.

Thus, the scrambled speech tokens had similar magnitude spectra and similar temporal-fine structure characteristics as the original speech utterances, giving them speech-like perceptual qualities. However, because the sequence of the acoustic snippets was shuffled, the scrambled speech was unintelligible.

Furthermore, the passive differed from the active task in that the handheld response vibrated randomly between 3 and 5 times during each block. Listeners were instructed to passively listen to the sounds and press the right trigger button on the handheld response interface each time the interface vibrated, ensuring that the listener stayed engaged in this task. Listeners need to correctly detect at least 2 out of 3 vibrations, otherwise they were excluded from the study.

In the active task of experiment 1, target and masker differed in both voice pitch and perceived spatial direction, and listeners could use either cue to direct their attention to the target voice. Experiment 2 further assessed the role of spatial attention in two active tasks. The first task (“spatial cues”) was identical to the active condition of Experiment 1. The second task (“no spatial cues”) used similar stimuli as the active task in experiment 1, except that both sources had 0 μs ITD. Thus, in experiment 2, each listener completed six blocks of an active listening task that was identical to the active task in experiment 1 and six blocks of another active listening task that was similar to the active task in experiment 1, except that the spatial cues were removed. Blocks were randomly interleaved. Listeners indicated when they detected the target talker uttering one of the four color words, by pressing the right trigger on the handheld response interface.

### Signal Processing of the fNIRS traces

We used HOMER2 (Huppert et al. 2009), a set of Matlab-based scripts, to analyze the raw recordings of the deep and shallow fNIRS channels at each of the 4 ROIs. First, the raw recordings were band-pass filtered between 0.01 and 0.3 Hz, using a fifth order zero-phase Butterworth filter. Next, we removed slow temporal drifts in the band-pass filtered traces by de-trending each trace with a 20th-degree polynomial (Pei et al., 2007). To remove artefacts due to sudden head movement during the recording, the detrended traces were then wavelet transformed using Daubechies 2 (db2) base functions. We removed wavelet coefficients that were outside of one interquartile range (IQR) (Molavi et al. 2012). We applied the modified Beer-Lambert law (Cope and Delpy, 1988; Kocsis et al., 2006) to these processed traces and obtained the estimated oxygenated hemoglobin (HbO) concentrations for the deep and shallow channels at each ROI. To partial out physiological nuisance signals, thus reducing across-listener variability, we then normalized all HbO traces from the task conditions by dividing them by the maximal HbO concentration change during controlled breathing.

### Calculation of Activation levels

For each of the auditory task conditions and ROIs, we wished to determine what portion of each hemodynamic response could be attributed to the behavioral task. Therefore, HbO traces were fitted by four general linear models (GLM), one GLM for each ROI. Each GLM was of the form:

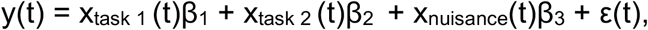

where y is the HbO trace, t is time, and the β_i_-values indicate the activation levels of each of the regressors. We calculated the β_i_-values for each listener and ROI. Specifically, x_task i_ (t) was the regressor of the hemodynamic change attributed to behavioral task i. x_nuiscance_(t) the HbO concentration in the shallow channel (Brigadoi and Cooper, 2015), and ε(t) the residual error of the GLM.

The task regressors x_task_ i in the GLM design matrix then contained reference functions for the corresponding task, each convolved with a canonical hemodynamic response function (HRF, Lindquist et al., 2009):

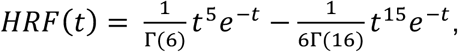

where Γ was the gamma function.

Task reference functions were built from unit step functions as follows. In the controlled breathing task, the reference function equaled 1 during the breath holding time intervals, and 0 otherwise. Only one task regressor was used to model the controlled breathing task. In the auditory tasks, two reference functions were built, one for each task, and set to 1 for stimulus present, and 0 for stimulus absent.

### Statistical Analysis

To assess whether the HbO activation levels at each ROI differed from 0, we applied two-sided Student’s t-tests. Furthermore, to determine whether HbO activation levels differed from each other across the two task conditions of each experiment, left/right hemispheres and dorsal (tgPCS)/ventral (cIFS) sites, 2×2×2 repeated-measures analyses of variance (rANOVA) were applied to the β_i_-values, at the 0.05 alpha level for significance. To correct for multiple comparisons, all reported p values were Bonferroni-corrected.

## RESULTS

### Controlled Breathing Task

Figure 3 shows the HbO traces during the controlled breathing task for both experiments 1 and 2, at each of the four ROIs. Two-sided Student’s t-test on the β- values of the GLM revealed that at each ROI, the mean activation levels during breath holding differed significantly from 0 [t(13) = −7, p < 0.001 at left tgPCS; t(13) = −7, p < 0.001 at right tgPCS; t(13) = −6.5, p < 0.001 at left cIFS; t(13) = −7.5, p < 0.001 at right cIFS, after Bonferroni corrections]. Two-sided Student’s t-test confirmed that also in experiment 2, HbO activation levels during breath holding significantly differed from 0 [t(13) = −5.6, p < 0.001 at left tgPCS; t(13) = −3.4, p < 0.001 at right tgPCS; t(13) = −4, p < 0.001 at left cIFS; t(13) = −3.7, p = 0.006 at right cIFS]. Thus, breath holding induced a significant change in the BOLD response at all four ROIs, confirming feasibility of the recording setup and providing a baseline reference for normalizing the task-evoked HbO traces of experiments 1 and 2.

**Figure 3.**
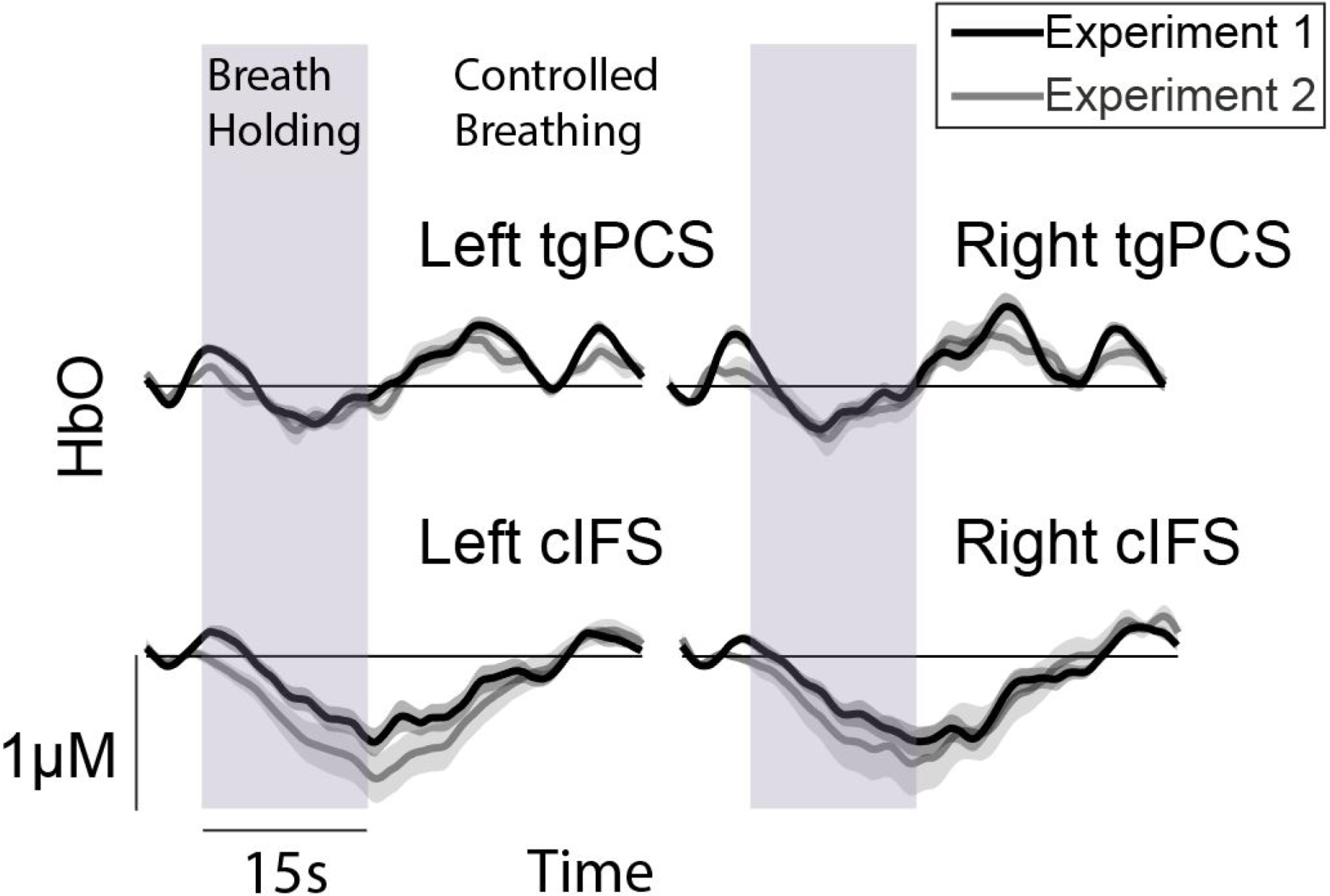
HbO concentration change during controlled breathing in experiments 1 and 2.

### Experiment 1

Figure 4A shows the HbO traces during active versus passive listening, at each of the four ROIs. Solid lines denote the auditory attention condition, dotted lines passive listening. The ribbons around each trace show one standard error of the mean across listeners. Figure 4B shows BOLD activation levels p, averaged across listeners, during the auditory attention (solid fill) and the passive listening tasks (hatched fill). Error bars show one standard error of the mean. All listeners reached criterion performance during behavioral testing and were included in the group analysis. RANOVA revealed significant main effects of task [F(1,13) = 6.5, p = 0.024] and dorsal (tgPCS)/ventral (cIFS) site [F(1,13) = 6.1, p = 0.028]. The effect of hemisphere was not significant [F(1, 13) = 0.015, p = 0.9]. In experiment 1, listeners were tested over 12 blocks, a number we initially chose conservatively.

**Figure 4.**
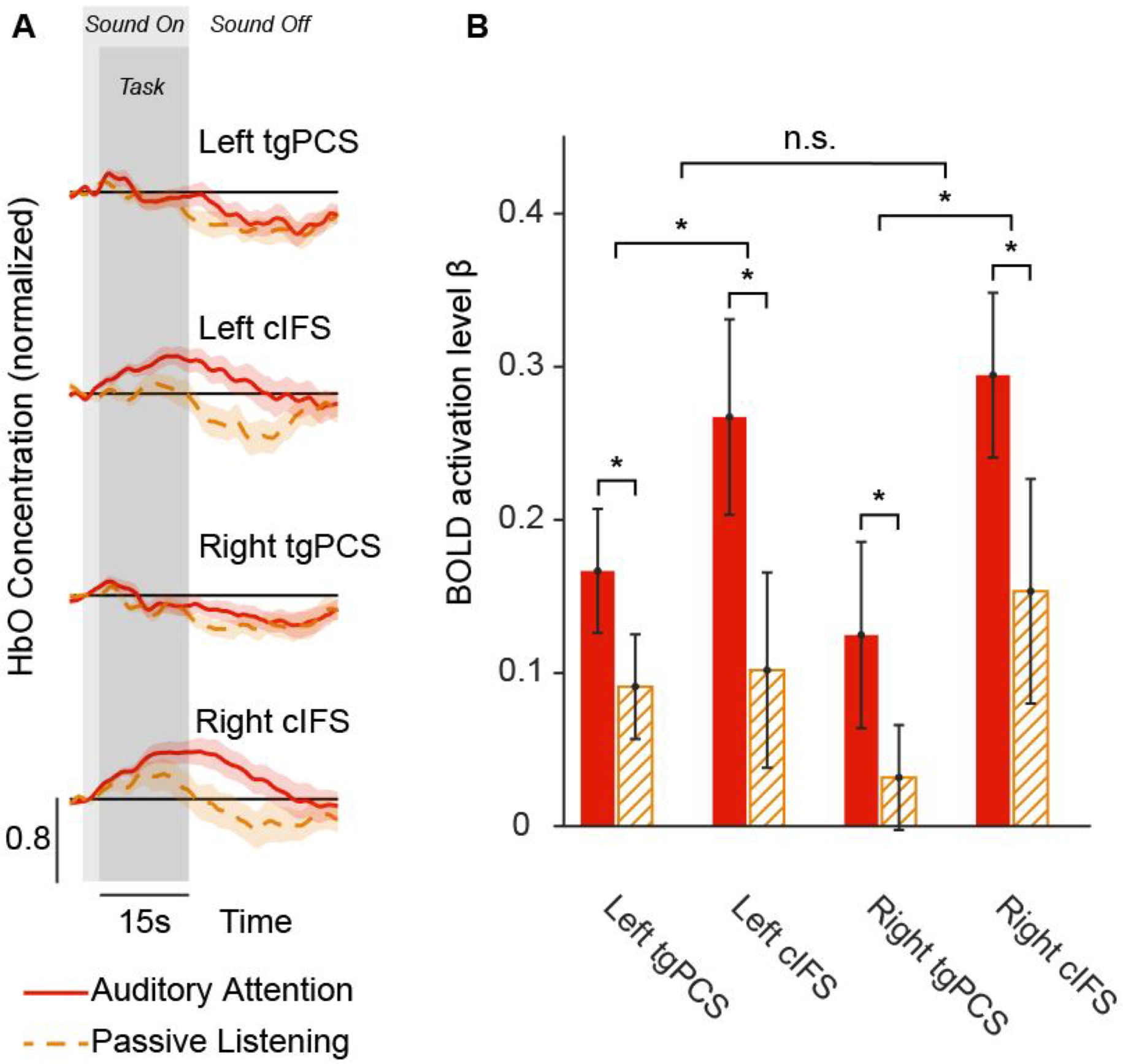
Results from experiment 1. A) Normalized HbO traces during the direction of auditory attention versus passive listening, at each of the four ROIs in experiment 1. The ribbons around each trace show one standard error of the mean across listeners. B) Normalized HbO traces during pitch and spatial cues condition versus pitch cue only condition, at each of the four ROIs in experiment 2. The ribbons around each trace show one standard error of the mean across listeners. BOLD activation levels β, error bars show one standard error of the mean.

To investigate the minimum number of blocks needed to see a robust difference between active and passive listening conditions, we applied a power analysis. Using bootstrapping of sampling without replacements, we calculated activation levels β during active versus passive listening in 100 repetitions and found that a minimum of 6 blocks suffices to show a robust effect. Therefore, in experiment 2, listeners were tested using 6 blocks per condition.

### Experiment 2

Figure. 5A and B display the HbO traces (red lines denote spatially separated, blue lines co-located configurations) and the across-listener average in BOLD activation β-levels for the spatially separated (red fill) versus co-located configurations (blue fill), at each of the four ROIs. 14 listeners reached criterion performance during behavioral testing and were included in the group analysis. One listener’s data had to be excluded, because the participant had fallen asleep during testing. An rANOVA on the activation levels found a significant main effect of dorsal/ventral site [F(1, 13) = 10.3, p = 0.007]. Main effects of spatial configuration and left/right hemisphere were not significant [F(1,13) = 1.6, p = 0.212 for effect of task; F(1,13) = 0.153, p = 0.702 for effect of hemisphere]. In addition, the interaction between task and left/right hemisphere was significant [F(1,13) = 7.2, p = 0.019], confirming an overall stronger activation in the right hemisphere in the spatially separated as compared to the co-located configuration.

**Figure 5.**
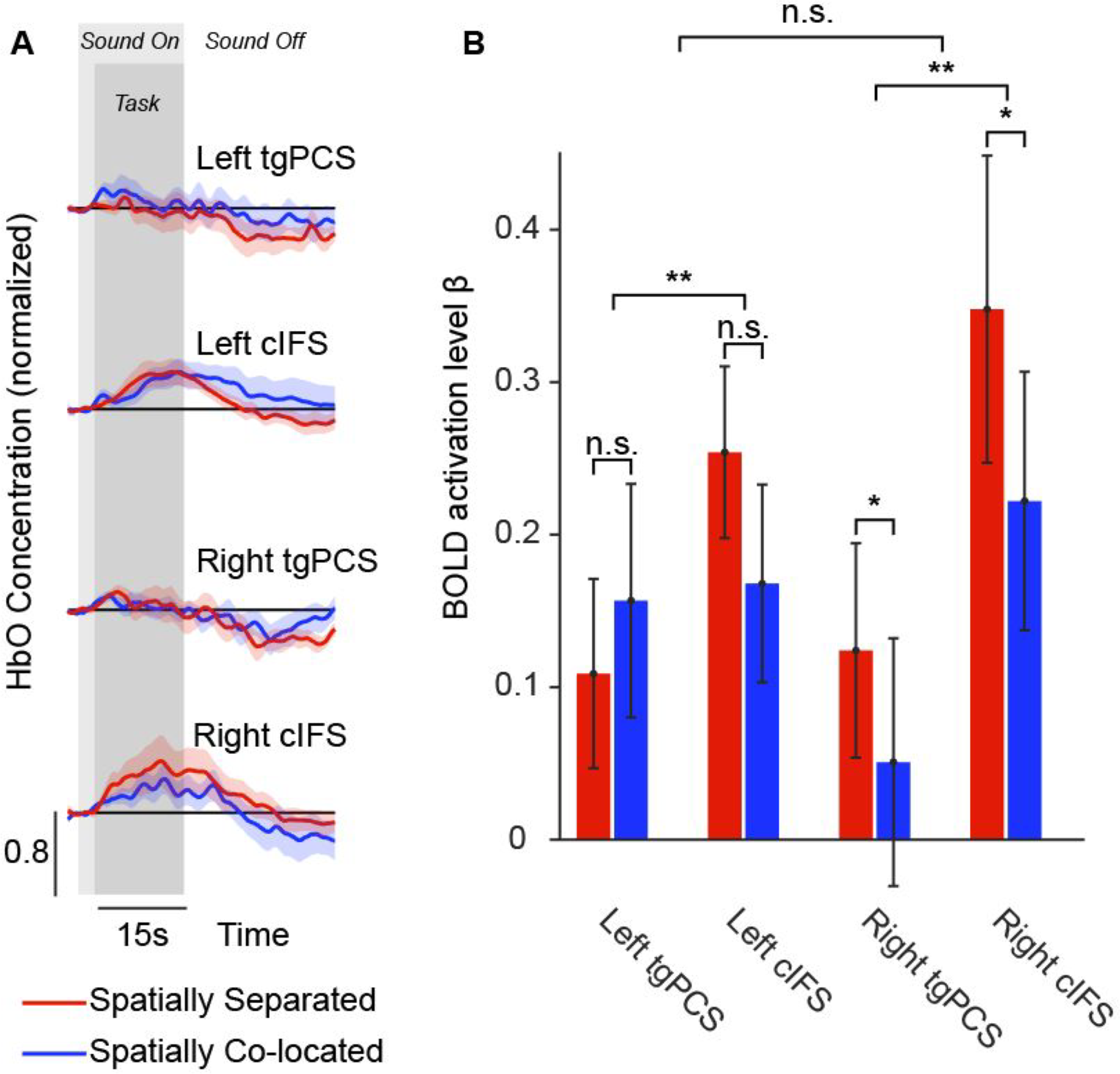
Results from experiment 2, formatting similar to Figure 4.

## DISCUSSION

### 1. Physiological correlates of active listening exist in LFCx

In experiment 1, we presented two competing streams of rapidly changing words. All target and masker words were drawn from an identical corpus of possible words, uttered by two male talkers and played synchronously. As a result, both EM and IM interfered with performance. When the sounds were unintelligible scrambled speech and the participants listened passively, across all ROIs, the LFCx responses were smaller as compared to the active auditory attention task. Thus, direction of auditory attention increased bilateral BOLD responses in LFCx. These results support and extend previous finding on the role of LFCx. Using rapid serial presentation task with two simultaneous talkers, where listeners monitored a target stream in search for targets and were tasked to detect-and-identify target digits, prior work had revealed an auditory bias of LFCx regions (Michalka et al., 2015). Here we found that even when listeners were performing a detection-only task under conditions of IM, this resulted in robust recruitment of LFCx. Moreover, the current results show that attentive listening in a situation with IM recruits LFCx, whereas passive listening does not.

### 2. Right LFCx activation associated with SRM

We wished to disentangle the role of spatial attention on the LFCx BOLD response. In experiment 1, spatial differences between target and masker were available. However, the target voice also had a slightly lower pitch than the masker voice, and listeners could utilize either or both cues to attend to the target (Ihlefeld and Shinn-Cunningham, 2008b). Therefore, we presented two different spatial configurations in experiment 2 - a spatially separated configuration, where spatial attention could help performance, and a spatially co-located configuration, where spatial attention cues were not available. Contrasting active listening across these two spatial configurations, experiment 2 revealed that right LFCx was more strongly recruited in the spatially separated as compared to the co-located configuration. In contrast, in left LFCx, no difference in BOLD signals was observed across the two spatial configurations. Therefore, these findings are consistent with the interpretation that right LFCx BOLD activation contained significant information about the direction of spatial attention.

In general, spatial release from masking is thought to arise from three different mechanisms (e.g., Shinn-Cunningham et al., 2005), monaural head shadow, assumed to be a purely acoustic phenomenon, binaural decorrelation processing, and spatial attention. The current stimuli did not provide head shadow. Therefore, in the current paradigm, spatial cues could have contributed to spatial release from masking through two mechanisms, binaural decorrelation, presumably arising at or downstream from the brainstem (Wong and Stapells, 2004; Dajani and Picton, 2006; Wack et al., 2012) and spatial attention, assumed to arise at cortical processing levels (Zatorre et al., 1999; Ahveninen et al., 2006; Shomstein and Yantis, 2006; Wu et al., 2007; Larson and Lee, 2014).

Alternatively, or in addition, a stronger BOLD response in the spatially separated versus co-located configurations could also be interpreted in support of the notion that right LFCx BOLD activity correlates with overall higher speech intelligibility in the spatially separated configuration. However, converging evidence from recent studies in NH listeners finds physiological correlates of speech intelligibility in the *left* hemisphere and at the level of auditory cortex as opposed to LFCx (Scott et al., 2009; Olds et al., 2016; Pollonini et al., 2014; Sheffield et al., *in press*). It is possible that here, listeners had to spend more listening effort in the spatially co-located versus separated configurations. However, comparing noise-vocoded versus unprocessed speech in quiet, or in competing background speech, previous work finds that increased effort differentially activates the *left* inferior frontal gyrus (Wiggins et al., 2016a; Wijayasiri et al., 2017). Moreover, testing NH listeners with a 2-back working memory task on auditory stimuli, Noyce and colleagues (2017) confirmed the existence of auditory-biased LFCx regions, suggesting that here, the observed physiological correlates of spatial release from masking may be caused by differences in utilization of short-term memory across the two spatial configurations. Together, the current findings support a hypothesis already proposed by others (Papesh et al., 2017) that a cortical representation of spatial release from masking exists, and suggest that assessment of right LFCx activity is a viable objective physiological measure of spatial release from masking.

Recent work shows that decoding of cortical responses is a feasible measure for determining which talker a listener attends to (e.g., Mesgarani and Chang, 2012; Choi et al., 2013; O’sullivan et al., 2014; Mirkovic et al., 2015). Moreoever, previous physiological work on speech perception in situations with EM or IM shows recruitment of frontal-parietal regions when listening to speech with EM (Scott et al., 2004) and suggests that the left superior temporal gyrus is differentially recruited for IM whereas recruitment of the right superior temporal gyrus is comparable for both types of masker (Scott et al., 2009). With the current paradigm, LFCx recruitment could be used to predict whether or not a listener attends to spatial attributes of sound, a question to be investigated by future work.

### 3. Utility of fNIRS as objective measure of auditory attention

A growing literature shows that fNIRS recordings are a promising tool for assessing the neurobiological basis of clinical outcomes in cochlear implant users (e.g., Dewey and Hartley, 2015; Lawler et al., 2015; McKay et al., 2016; van de Rijt, et al., 2016). Cochlear implants are ferromagnetic devices, and when imaged with Magnetic Resonance Imaging (MRI), electroencephalography (EEG), or magnetoencephalography (MEG), the implants typically cause large electromagnetic artifacts and are sometimes even unsafe for use inside the imaging device. In contrast to MRI, EEG and MEG, fNIRS uses light to measure BOLD signals and thus does not produce electromagnetic artifacts when used in conjunction with cochlear implants. Moreover, compared to fMRI machines, fNIRS scanners are quiet, they do not require the listener to remain motionless and are thus more child-friendly (c.f., Bortfeld et al., 2009), and they are generally more cost effective.

However, previous work using fNIRS for assessing auditory functions found highly variable responses to auditory speech at the group level (Wiggins et al., 2016b). To reduce across-listener variability, here, we used the individual’s own maximal amplitude during controlled breathing for normalizing the HbO traces during the auditory task, followed by fitting a GLM where we regressed out nuisance signals from a shallow trace that recorded blood oxygenation close to the surface of the skull. Results demonstrate that fNIRS is a feasible approach for characterizing central auditory function in NH listeners.

Objective measures of masked speech identification in IM could, for instance, be used to assess the neurobiological basis for predicting rehabilitative success in newly implanted individuals. A long-term goal of our work is thus to establish an objective measure of auditory attention that could be used to study central nervous function in cochlear implant users. Here we find that fNIRS is a promising tool for recording objective measures of spatial auditory attention in NH listeners, with potential application in cochlear implant users.

### 4. Conclusions

Two experiments demonstrated that when NH listeners are tasked with detecting the presence of target keywords in a situation with IM, bilateral LFCx BOLD responses, as assessed through fNIRS, carry information about whether or not a listener is attending to sound. In addition, right LFCx responses were stronger in a spatially separated as compared to a co-located configuration, suggesting that right LFCx activity is associated with spatially directed attention.

## ACKNOWLEDGMENTS

This work was supported by the New Jersey Health Foundation (PC 24-18 to AI) and the National Science Foundation (MRI CBET 1428425 to T Alvarez and B Biswal).

